# Arabidopsis cell suspension culture and RNA sequencing reveal regulatory networks underlying plant programmed cell death

**DOI:** 10.1101/2023.03.14.532467

**Authors:** Rory Burke, Aideen McCabe, Neetu Ramesh Sonawane, Meet Hasmukh Rathod, Conor Whelan, Paul F. McCabe, Joanna Kacprzyk

**Affiliations:** School of Biology and Environmental Science, University College Dublin, Dublin 4, Ireland

**Author notes:** Corresponding author email: Dr. Joanna Kacprzyk The author(s) responsible for distribution of materials integral to the findings presented in this article in accordance with the policy described in the Instructions for Authors is: Dr. Joanna Kacprzyk.

**Keywords:** Plant Programmed Cell Death, *Arabidopsis thaliana*, Cell suspension culture, transcriptomics, salicylic acid, heat response

## Abstract

Programmed cell death (PCD) facilitates targeted elimination of redundant, damaged, or infected cells via genetically controlled pathways. In plants, PCD is often an essential component of normal development and can also mediate responses to abiotic and biotic stress stimuli. However, studying the transcriptional regulation of this fundamental process is hindered by difficulties in sampling small groups of cells undergoing PCD that are often buried within the bulk of living plant tissue. We addressed this challenge by using RNA sequencing (RNA-Seq) of *Arabidopsis thaliana* suspension cells, a system that allows precise monitoring of PCD activation and progression. The use of three PCD-inducing treatments (salicylic acid, heat and critical dilution), in combination with three cell death modulators (3- methyladenine, lanthanum chloride and conditioned medium), allowed isolation of candidate ‘core’ and stimuli-specific PCD genes, inference of underlying gene regulatory networks and identification of putative transcriptional regulators. This analysis underscored cell cycle disturbance and the repression of both pro-survival stress responses and mitochondrial retrograde signalling as key elements of the PCD-associated transcriptional signature in plants. Further, phenotyping of twenty *Arabidopsis* T-DNA insertion mutants in selected candidate genes confirmed a role for several in PCD and stress tolerance regulation, and validated the potential of these generated resources to identify novel genes involved in plant PCD pathways and/or stress tolerance in plants.

## Introduction

Programmed cell death (PCD) is a genetically regulated pathway for selective elimination of unwanted or damaged cells (Ameisen, 2002). In plants, PCD is an inherent component of normal development (Daneva et al., 2016), and may occur in response to abiotic and biotic stresses (Petrov et al., 2015; Coll et al., 2011). For example, localized PCD events can limit the spread of invading biotrophic pathogens (Balint-Kurti, 2019), mediate cold acclimation (Hong et al., 2017) and drive anatomical adaptations to salinity (Huh et al., 2002) and hypoxia (Evans, 2004). As the frequency and intensity of environmental stresses that negatively impact plant growth and productivity increases (Zandalinas et al., 2021), a greater understanding of PCD regulation is urgently required to inform the development of crops capable of withstanding future climatic conditions.

In contrast to the multiple forms of metazoan PCD that have been extensively characterised (Galluzzi et al., 2018), our understanding of cell death regulatory pathways in plants is limited. While there are similarities between PCD programmes operating in plants and animals (Rantong and Gunawardena, 2015), the homologues of core regulators of PCD in animal models, such as caspases and BCL-2 family proteins, have not been identified in plant genomes (Sueldo and van der Hoorn, 2017; Dickman et al., 2017). Decades of intense research has uncovered many plant-specific elements of the PCD pathway, including the BAG (Bcl-2 associated athanogene) protein family (Thanthrige et al., 2020) as well as an array of proteases driving cell death execution (Balakireva and Zamyatnin, 2019). Nevertheless, the sequence of events leading to PCD initiation and execution in plants, as well as the underlying gene regulatory networks (GRNs), requires further elucidation.

The core PCD machinery in animals is regulated at post-translational level, largely through changes in protein localization and proteolytic cascades (Fuchs and Steller, 2015; Cubría-Radío and Nowack, 2019). However, gene regulation also contributes to PCD initiation and execution, with the expression of many pro-apoptotic genes controlled by transcription factors (TFs) such as the tumour suppressor p53 (Aubrey et al., 2018). PCD regulation at the level of gene expression is also likely in plants, as the mRNA synthesis inhibitor actinomycin D and the translation inhibitor cycloheximide have both been shown to modulate PCD rates in cultured plant cells (Vacca et al., 2004; Masuda et al., 2003); Doyle et al., 2010). Indeed, numerous TFs implicated in both developmental (Cubría-Radío and Nowack, 2019) and stress-induced (Burke et al., 2020) PCD have been identified. Characterization of PCD-associated transcriptional signatures is therefore a promising strategy to advance our understanding of this important process.

To date, several studies have been undertaken to characterize the transcriptional changes occurring during plant PCD. Meta-analysis of microarray datasets associated with a variety of accepted or hypothetical PCD contexts has successfully isolated several ‘core’ genes associated with developmental cell death (Olvera-Carrillo et al., 2015). Further, RNA-Seq has been used to examine changes in gene expression during different forms of developmental PCD (Reza et al., 2018), (Li et al., 2019), immune cell death induced by pathogens or effectors (Matuszkiewicz et al., 2018; Bahieldin et al., 2016) and PCD in response to abiotic stressors such as boron deprivation (Kobayashi et al., 2018) or ionizing radiation (Johnson et al., 2018). However, many of these datasets were generated using RNA isolated from whole organs, as sampling only the cells undergoing PCD, often buried in the bulk of the tissue, can be technically challenging. Another drawback of the currently available datasets is the lack of quantitative determination of PCD rates in the sequenced samples, or even evidence that PCD was indeed occurring. As such, many studies that aim to describe transcriptional signatures of cell death suffer from low specificity for PCD rather than uncontrolled tissue damage or sub-lethal levels of stress, and a lack of spatial and temporal precision to enable only cells undergoing PCD to be sampled at the optimum time point. As a result, distinguishing transcriptional regulators of PCD from genes controlling processes preceding or counteracting cell death, such as stress acclimation, can be challenging. Finally, studies using a single elicitor do not allow identification of genes associated with PCD from those linked to a general response to stress or associated with the response to the specific elicitor, and thus may struggle to definitively identify *bona fide* PCD regulators. To overcome these challenges, specific transcriptome profiles of well-established PCD systems should be generated to achieve a higher resolution both in terms of PCD quantification and identification of differentially regulated genes (Olvera-Carillo et al 2015).

We accomplished that in this study by using *Arabidopsis thaliana* cell suspension culture (ACSC), a widely used model for studying plant PCD (Malerba and Cerana, 2021). The ACSC consists of a homogenous cell population in which cell death can easily be induced by a variety of chemical or physical stimuli (McCabe and Leaver, 2000; Lanubile et al., 2022). Further, the progression of cell death can be monitored using phase-contrast microscopy that also enables differentiation between PCD and uncontrolled necrotic cell death based on the hallmark PCD morphology: protoplast condensation and retraction away from the cell wall (Reape et al., 2008; Kacprzyk et al., 2017). Finally, the ACSC facilitates the application of chemical modulators of PCD pathways that may delay viability loss or shift cell fate towards uncontrolled cell death (necrosis) (McCabe et al., 1997; Smith et al., 2015; Kacprzyk et al. 2017; Awwad et al., 2019). The ACSC system thus offers a unique opportunity to specifically sample plant cells undergoing PCD and to identify the associated changes in gene expression. Indeed, it has been used previously for microarray-based analyses of heat-induced and senescence-related PCD (Swidzinski et al., 2002) and cell death activated by the herbicide Rose Bengal (Gutiérrez et al., 2014).

In this study, we used 3 different PCD inducers (salicylic acid, SA; heat stress, HS; critical culture dilution, CD) and 3 different PCD inhibitors (3-methyladenine, 3- MA; lanthanum chloride, LaCl_3_; conditioned medium, CM). Comparing the transcriptional responses induced in the ACSC by different stimuli (PCD inducers and PCD modulators) enabled changes in gene expression related to PCD pathway itself to be distinguished from those that are specific to a particular treatment. We generated unique RNA-Seq datasets that provided a snapshot of gene expression changes occurring in cells committed to PCD. This allowed for analysis of upstream transcriptional regulators, as well as inference of ‘core’- and stimuli-specific GRNs. Subsequent phenotyping of Arabidopsis T-DNA insertion mutants functionally validated the role of selected candidate genes in PCD regulation and stress responses, further expanding our understanding of these fundamental processes in plants and underscoring the value of the generated resources.

## Results

### Induction and modulation of PCD in *Arabidopsis thaliana* Suspension Cells

We used three PCD-inducing treatments to gain insights into the transcriptional regulation of cell death events induced by different types of stimuli in *Arabidopsis thaliana* (Fig 1A). SA is a phytohormone implicated in regulation of PCD triggered in response to pathogens (Alvarez, 2000; Brodersen et al., 2005). HS induces oxidative stress dependent PCD (Vacca et al., 2004), and CD disrupts social signaling through dilution of survival signals the suspension cells release to the growth medium to suppress the default death programme (McCabe et al 1997). Applied treatments thus mimicked PCD induced by biotic interactions, abiotic stress and developmental programmes in the ACSC system. Further, to capture changes in gene expression associated with the PCD pathway, rather than a general response to each cell death stimuli, we used treatments previously reported to inhibit PCD in plant models (Fig 1A). 3-MA, an autophagy (Takatsuka et al., 2004; Kosic et al., 2021) and mitochondrial permeability transition blocker (Xue et al., 2002), was used to inhibit PCD induced by SA (Kacprzyk et al., 2014). LaCl_3_, an extracellular calcium channel blocker (Lewis and Spalding, 1998), was applied to inhibit PCD triggered by heat (Kacprzyk et al., 2017). CM, containing survival signals released by the suspension cells over 6-day period, was employed to decrease PCD rates resulting from CD (McCabe et al 1997).

**Figure 1.**
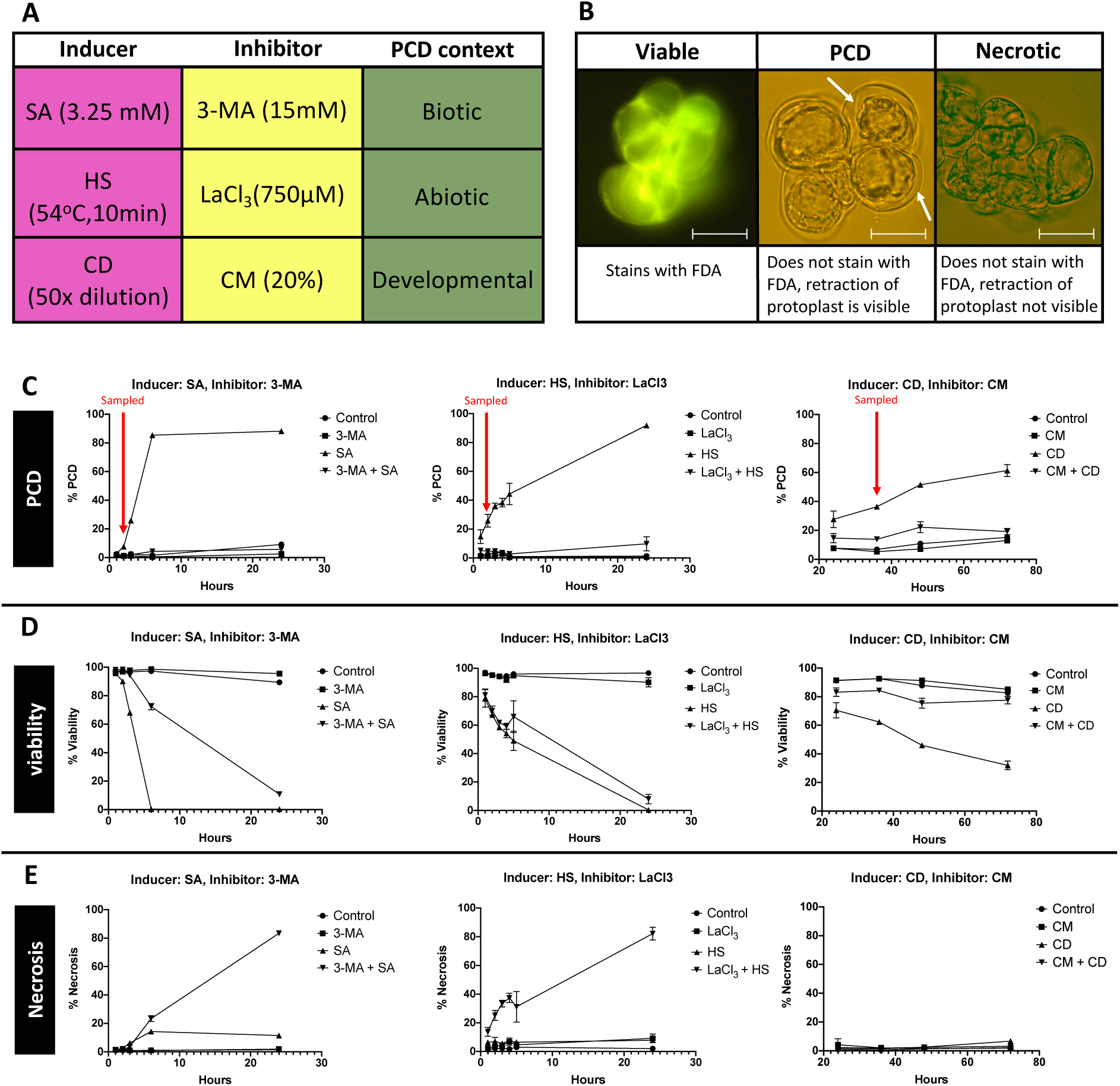
Response of ACSC to PCD inducer/inhibitor combinations. Summary of combinations of PCD inducers/inhibitors **(A).** Representative morphology of viable, PCD and necrotic ACSC cells stained with FDA. White arrows indicate the protoplast retraction characteristic of PCD, scale bar = 40 µm **(B)**. Time course data for PCD **(C)**, viability **(D)** and necrosis rates induced in ACSC by treatment with SA, w/wo 24h pretreatment with 3-MA; HS, w/wo 10 min pretreatment with LaCl_3_ and CD, w/wo CM added. The time points at which cells were sampled for RNA profiling are indicated by the red arrows. Experiments were repeated 3 times, and error bars display SEM. Where not visible, SEM is within the area covered by the symbol.

The effect of applied treatments on the rates of PCD, necrosis, and viability of cultured cells was determined as previously described (Reape et al., 2008), based on fluorescein diacetate (FDA) staining and presence of protoplast shrinkage morphology (Fig 1B). The levels of the PCD inducing treatments (i.e. concentration of SA, temperature/duration of HS, degree of cells dilution) were optimized to achieve high rates of PCD, but not necrosis; while the concentrations of 3-MA, LaCl_3_ and CM used resulted in significant PCD inhibition, without affecting viability rates in control cells (Fig 1 CDE). The PCD response induced by each treatment was monitored over the course of 1-24 h for SA and HS experiments, and 24-72 h for CD experiment to identify a time point when PCD rates have started to increase, but before the PCD morphology could be visually detected in the majority of the treated cells. At these time points (SA: 2h, HS: 2h, CD: 36h) cells were harvested for transcriptomic profiling to characterize changes in gene expression occurring during PCD induction and at early stages of PCD execution. The 3-MA and LaCl_3_ treatments resulted in significant inhibition of PCD levels induced by SA and HS respectively, with cells exhibiting delayed viability loss followed by necrosis; whereas the CM-treated cells cultured at low density maintained high viability throughout the 72-hour monitoring period. The significantly lower rates of PCD induced by SA, HS and CD in the presence of 3-MA, LaCl_3_ and CM respectively (Fig 1C) confirmed that our experimental setup offers a promising opportunity to draw a distinction between changes in gene expression that are related to PCD, and those specific to the stress treatment applied or to a general stress response.

### Transcriptional response associated with PCD induction in *Arabidopsis* suspension cells by range of stimuli

For each PCD inducer/blocker combination (SA/3-MA, HS/LaCl3, CD/CM), we identified genes differentially expressed in samples showing the PCD phenotype (“PCD”) compared to samples displaying low rates of PCD (“no-PCD”). This involved isolating (i) genes differentially expressed in samples treated with the PCD inducer (“PCD”) compared to control (‘no-PCD’) (SA vs control, HS vs control, CD vs control); and (ii) genes differentially expressed in samples treated with the PCD inducer (“PCD”) compared to samples treated with the PCD inducer in presence of the PCD inhibitor (“no-PCD”) (SA vs 3-MA+SA, HS vs HS+LaCl_3_, CD vs CD+CM) (Fig 2A). The effect of applied PCD blockers alone was also investigated (3-MA vs control, LaCl_3_ vs control, CM vs control) (Fig 2A). The vast majority of differentially expressed genes (DEGs) at 0.05 adjusted *p*-value cut-off had the log_2_ Fold Change (LFC) >0.25 (Fig 2A). The full DESeq2 results for the comparisons described above are provided in Table S1. These analyses highlighted extensive transcriptional changes induced by SA and HS, compared to transcriptional response to dilution below the critical cell density (CD) at the investigated time points (Fig 2A).

**Figure 2.**
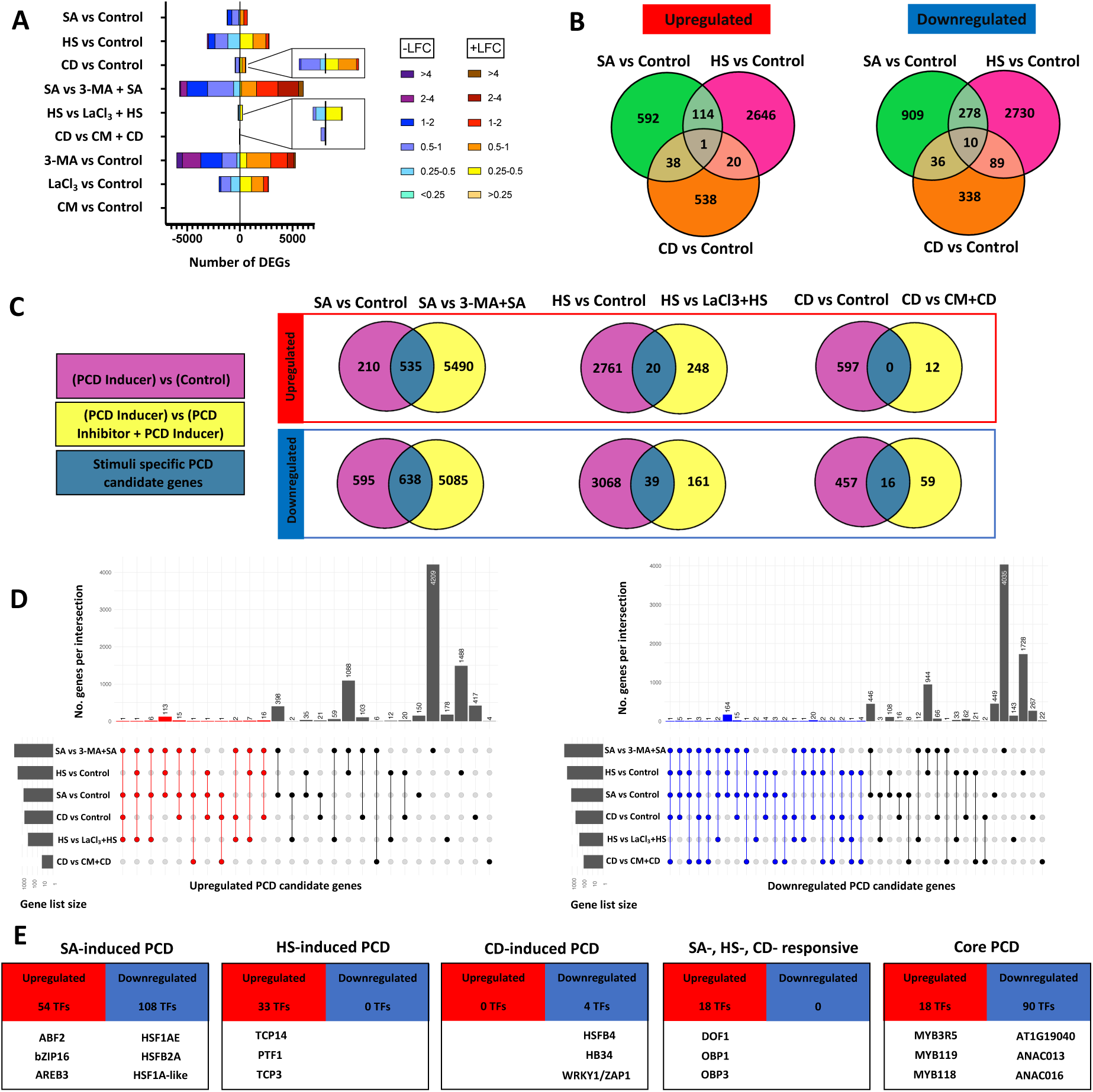
Transcriptional response to applied PCD inducers and PCD inhibitors. Number of DEGs and their Log_2_ Fold Change (LFC) distribution for indicated treatment comparisons (“PCD vs no-PCD”). The differential gene expression (DGE) analysis was performed for samples showing PCD phenotype (i.e. PCD inducer treatment: SA, HS, CD) compared to samples with low rates of PCD: (i) control samples, (ii) samples where PCD inducer was applied in combination with PCD inhibitor (i.e. 3-MA+SA, LaCl3+HS, CM+CD). The effect of PCD inhibitors alone (3-MA, LaCl3, CM) is also highlighted **(A)**. Venn diagrams presenting intersection between DEGs responsive to three used stress treatments (SA, HS, CD) **(B).** Venn diagrams presenting intersection between DEGs identified by conducted “PCD vs no-PCD” comparisons for each PCD inducer/inhibitor combination (SA/3-MA, HS/LaCl_3_, CD/CM) **(C).** UpSet plots present the overlapping upregulated (red) and downregulated (blue) DEGs in six “PCD vs no-PCD” comparisons performed for all PCD inducer/inhibitor combinations **(D)**. Set size represents number of DEGs for each “PCD vs no-PCD” comparison and intersection size corresponds to the number of shared DEGs among the comparisons marked as connected dots below. The single dots mark DEGs unique to respective “PCD vs no-PCD” comparisons. Upstream transcriptional regulators of the stimuli- and ‘core’- PCD candidate genes, identified using PlantTFDB TF Enrichement **(E).**

The differences in global gene expression of samples treated by PCD inducers in the presence or absence of PCD inhibitors were the largest in case of SA and 3-MA pre-treatment (SA vs 3-MA+SA: 11748 DEGs), compared to 468 DEGs for HS vs LaCl_3_+HS comparison, and only 87 genes differentially expressed between cells diluted below critical density with or without the addition of CM (Fig 2A). The effect of treatment with CM alone was minimal compared to the broad transcriptional shifts associated with treatment with 3-MA or LaCl_3_. The Metascape platform (Zhou et al., 2019) was used for enrichment analysis of genes responsive to 3-MA and LaCl_3_. The top 300 most significantly differentially regulated genes (by adjusted p-value) were used as an input, as Metascape recommends narrowing the gene list to this number to get meaningful results. The terms representing enriched clusters are presented in Fig S1, suggesting that both PCD modulators regulated pathways related to various aspects of stress responses and signalling. Moreover, as 3-MA was previously reported to exert autophagy independent effect on plant mitochondria (Kacprzyk et al., 2014), and mitochondria have a well-established role in the regulation and/execution of PCD process in plants (Van Aken and Van Breusegem, 2015), we inspected the effect of 3-MA on the expression of genes encoded by the mitochondrial genome and identified downregulation of all detected mitochondrial genes after incubation with 3-MA, that was not observed after treatment with LaCl_3_.

Comparison of the transcriptional response to the three used stress treatments (SA, HS, CD) revealed just 1 upregulated and 10 downregulated genes in common (Fig 2B, listed table S2). Subsequently, the candidate genes involved in regulation of PCD induced by each stress stimuli (SA, HS, CD) were shortlisted by selecting DEGs identified by both “PCD vs no-PCD” comparisons performed for each experiment (i.e. “PCD inducer” vs “Control”, and “PCD inducer” vs. “PCD inhibitor + PCD inducer”) and showing the same direction of regulation (i.e. either upregulated in both comparisons, or downregulated in both comparisons) (Fig 2C). Using this approach, sets of 1173, 59 and 15 candidate PCD genes were identified for SA-, HS- and CD-treatments respectively (listed Table S3). The candidates for stimuli independent, ‘core’ PCD genes, were shortlisted by selecting DEGs identified by at least half of all performed “PCD vs no-PCD” comparisons (SA vs control, HS vs control, CD vs control, SA vs 3-MA+SA, HS vs HS+LaCl3, CD vs CD+CM) and showing the same direction of regulation (Fig 2DE) yielding a list of 390 known genes (401 including novel transcripts: 164 upregulated and 237 downregulated) (listed Table S4). The Gene Ontology (GO) resource was used to analyse enrichment for biological processes among the 401 ‘core’ transcripts (listed Table S4). To ascertain the upstream transcriptional regulators of the shortlisted stress stimuli- and ‘core’- PCD candidate genes, the PlantTFDB ‘TF Enrichment’ tool (Jin et al., 2017) was used to find TFs with overrepresented targets (Table S5). The number of identified TFs and top 3 most significantly enriched TFs for each gene list are shown in Fig 2E.

### Construction of Regulatory Networks

To generate pathway level insights into observed transcriptional responses, and to further uncover the mechanisms underlying PCD, clustered GRNs were constructed using candidate gene lists isolated for stimuli specific (Fig 3) and ‘core’ (Fig 4) PCD programmes. The constructed GRNs facilitate identification of highly connected, hub signaling genes, that are likely to play a central role in the represented PCD pathway. Gene enrichment analysis was carried out on identified clusters within each network, highlighting biological processes relevant to stimuli specific and ‘core’ PCD programmes operating in ACSC.

**Figure 3:**
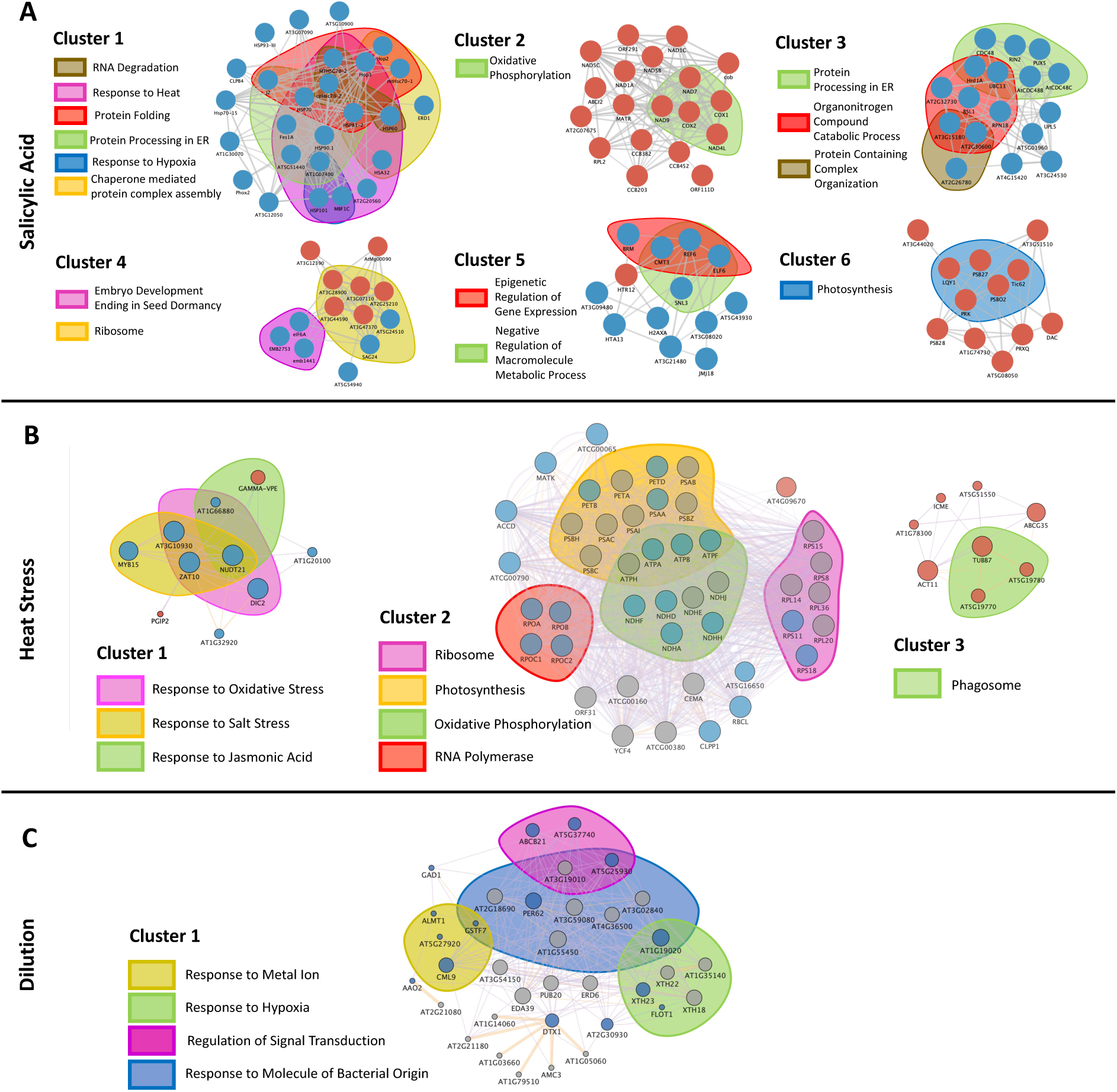
Stress stimuli specific GRNs underlying PCD. Clustered GRNs constructed in Cytoscape using candidate lists of stimuli specific PCD candidate genes. Red nodes represent genes upregulated and blue nodes downregulated under PCD-inducing conditions. **(A)** GRN for PCD induced by SA (and inhibited by 3-MA) where edges represent highly confident (confidence score = 0.7) protein-protein interactions provided by the String protein query database. **(B)** GRN for PCD induced by HS (and inhibited by LaCl_3_). **(C)** GRN for PCD induced by CD and inhibited by CM. In **B** and **C,** ‘Result’ nodes inserted by GeneMANIA are colored in grey, with node size representing the connectivity of each node (Degree score value, with more highly connected nodes appearing larger). Purple edges represent co-expressed genes, yellow edges represent predicted regulatory interactions and blue edges represent co-localized genes. Edge thickness represents normalized max weight, i.e. predicted strength of the interaction. Gene enrichment was carried out separately for each cluster using Metascape, and genes highlighted based on the significant gene enrichment terms to which they contribute.

**Figure 4.**
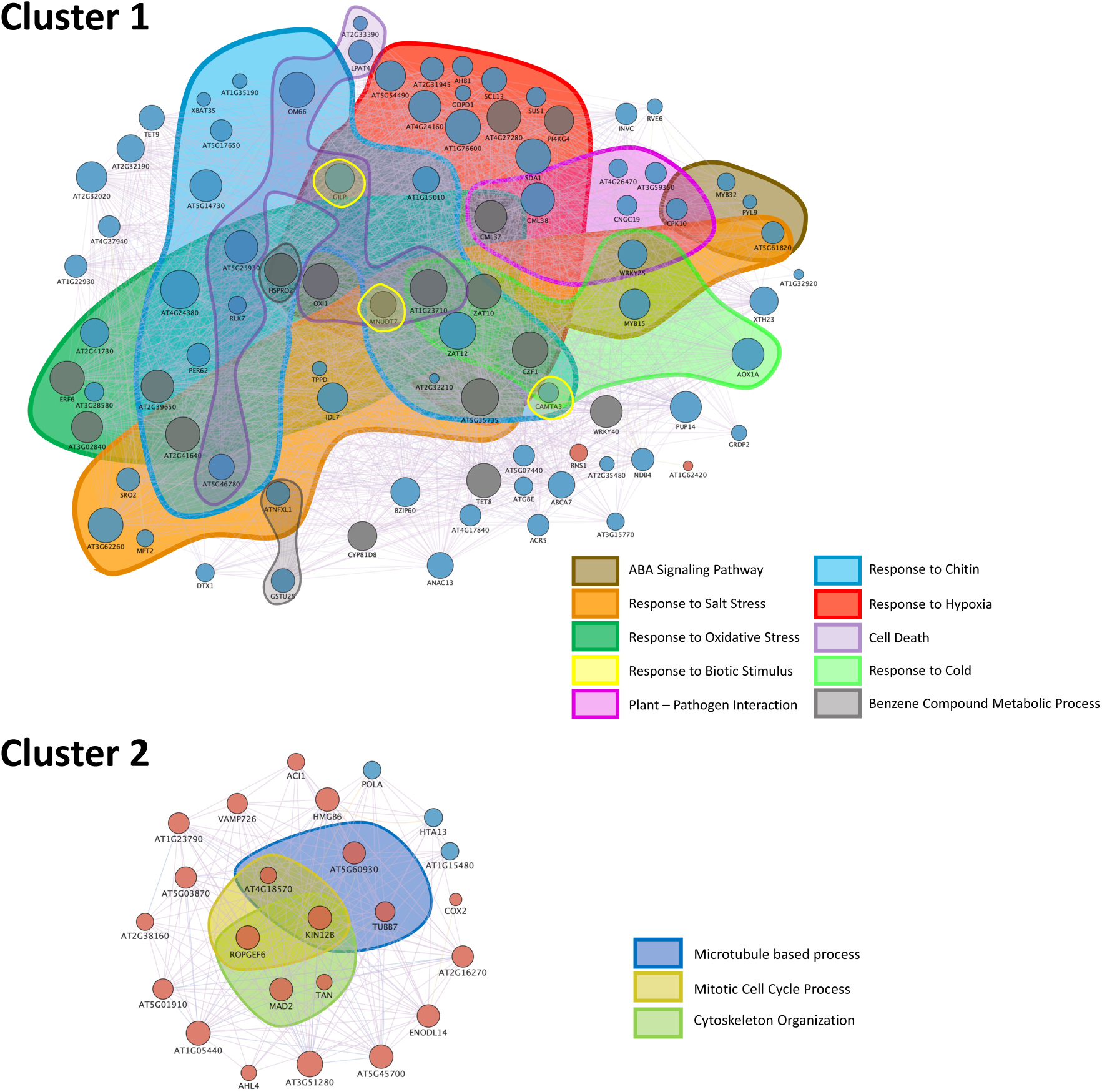
GRN underlying the ‘core’ PCD response. The clustered GRN was constructed using GeneMANIA using 390 ‘core’ candidate PCD genes identified from DGE analyses (listed Table S7). Red nodes represent genes upregulated and blue nodes downregulated under PCD-inducing conditions. Result’ nodes inserted by GeneMANIA are colored in grey, with node size representing the connectivity of each node (Degree score value, with more highly connected nodes appearing larger). Purple edges represent co-expressed genes, yellow edges represent predicted regulatory interactions and blue edges represent co-localized genes. ‘Result’ nodes inserted by GeneMANIA appear grey, with node size representing the connectivity of each node (degree score value, with more highly connected nodes appearing larger). Purple edges represent co-expressed genes, yellow edges represent predicted regulatory interactions and blue edges represent co-localized genes. Edge thickness represents normalized max weight, ie predicted strength of the interaction. Gene enrichment was carried out separately for each cluster using Metascape, and genes highlighted based on the significant gene enrichment terms to which they contribute.

### Phenotypic validation of selected PCD candidate genes

In order to investigate the potential of the generated transcriptomic resources to accelerate the discovery of new genes involved in plant PCD regulation, we shortlisted 20 genes for PCD and stress resilience phenotyping experiments. As detailed in Table S6 the selected genes were (i) highlighted as potential PCD regulators by RNA-Seq analyses performed herein, (ii) their involvement in plant PCD was evaluated as likely based on literature research, and (iii) there was homozygous, single T-DNA insertion *Arabidopsis thaliana* mutant lines available from the Nottingham Arabidopsis Stock Centre (NASC). Lines with a T-DNA insertion reported in the exon or coding sequence (CDS), were preferentially selected (Table S6). In order to evaluate the PCD response of selected T-DNA insertion mutants, the root hair assay, a method based on examination of viability and morphology of root hair cells (Kacprzyk et al., 2014) was used as it provides a rapid method for quantitative determination of PCD in mutant genotypes. Heat stress (HS; 45°C, 10 min) and 30 μM SA were used to induce PCD in *Arabidopsis thaliana* root hairs and rates of PCD were scored 6 and 24 hours after HS treatment, and 24 hours after SA treatment (Kacprzyk et al., 2016) Three rounds of screening were performed on all 20 mutant lines; and lines with an altered stress-induced PCD phenotype in root hairs were identified. Subsequently, the root hair assay repeated a further 2 times on each line to confirm the phenotype. Two mutant lines (*MYB3R-4* and *GAPC1*) displayed severely reduced germination which prevented phenotyping of these plants.

A reduced PCD phenotype in response to SA and/or HS treatment compared to wild type (Col-0) was observed for T-DNA insertion lines in *DTX-1* (SALKseq_064435.5), *MYB119-1* (SALK_015263C), *CMT3* (SALK_148381C) and *PGIP2* (SALK_014561C) (Fig 5A). In all cases, the reduction in PCD response was associated with an increase in viability, rather than a shift towards necrotic cell death (Figure S2). There is well established evidence linking PCD to seed development and germination (Matilla, 2019), and knockouts of PCD-associated genes has previously been shown to affect germination rates under abiotic stress in *Arabidopsis* (Bahieldin et al., 2016). Therefore, to further examine the potential link between *DTX-1*, *MYB119*, *CMT3* and *PGIP2* genes and PCD, the germination rates of each T-DNA insertion line was examined under control and stress conditions: NaCl, the herbicide methyl viologen (MV), the plant hormone SA, and the heavy metal cadmium (Cd) (Fig 5B). The *dtx1* plants displayed reduced germination compared to Col-0 on 0.5 μM MV, while the *cmt3* plants demonstrated increased germination on plates containing 2 μM MV. The *myb119* line showed reduced germination rate relative to Col-0 under control and all stress conditions, but did not appear to be more sensitive to stress than Col-0 as highlighted by its germination under stress conditions relative to control conditions (Figure S2). The stress sensitivity of *dtx-1*, *myb119*, *cmt3* and *pgip2* plants was also examined by transferring 7-day old seedlings to plates containing NaCl, SA or MV and determining survival rates after 7 days (Fig 5C). This identified higher survival rates of *dtx1* plants relative to Col-0 on 180mM NaCl, 400 μM SA and 2 μM MV, and increased survival of *myb119* plants on NaCl and SA treated plates.

**Figure 5.**
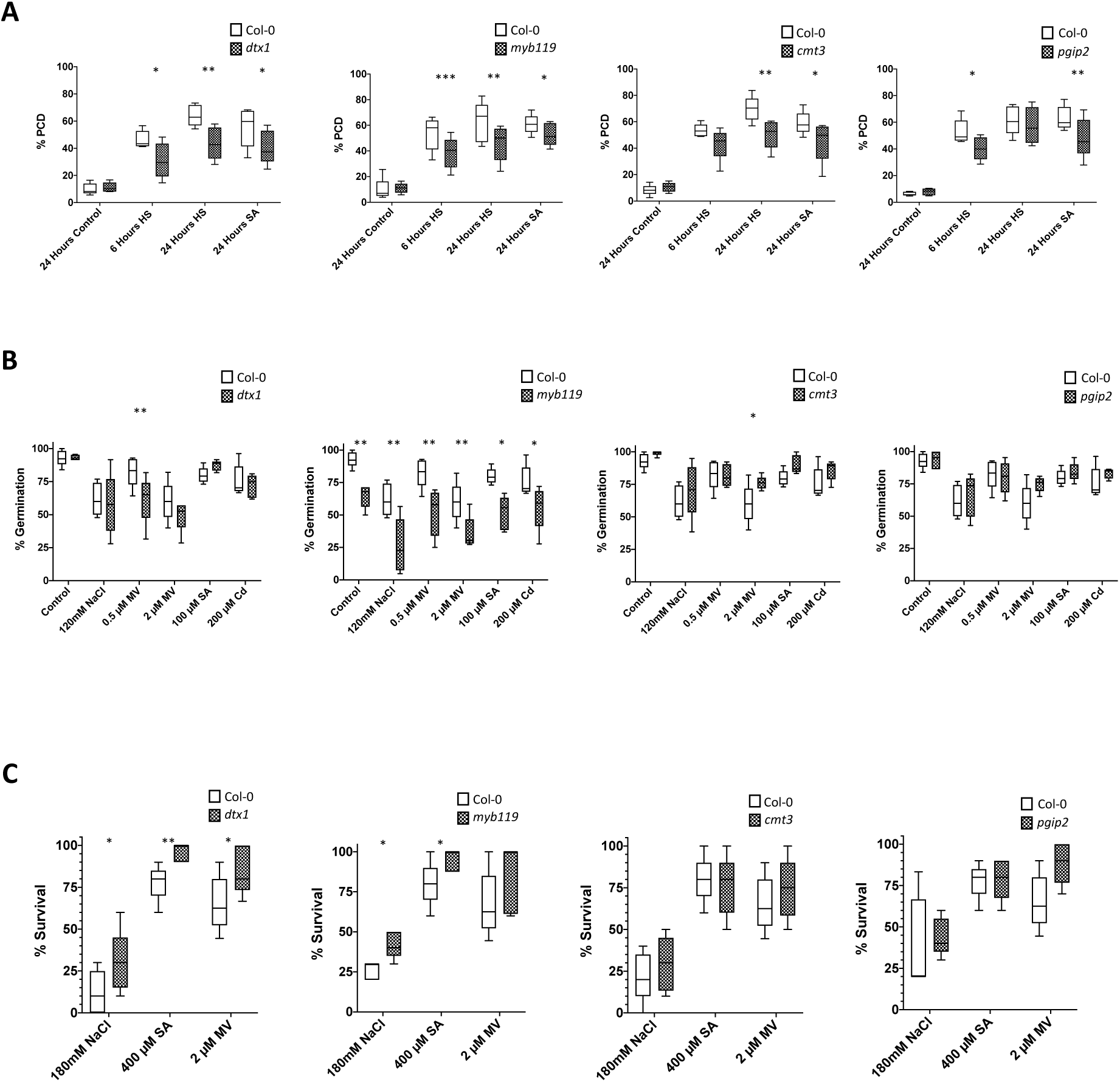
PCD and stress response phenotypes of *dtx-1, myb119, cmt3* and *pgip2* T-DNA insertion mutants. (A) PCD rates in seedling root hairs. 5-day old seedlings were transferred to a 24 well plate, with each well containing 1 mL of SDW. Seedlings were then subjected to either a 10-minute HS at 45°C or treated with 30 μM SA and scored for rates of PCD, viability and necrosis in root hairs after 6 and 24 hours (HS) or 24 hours (SA). Root hairs of 2 seedlings/genotype/experiment were examined, and experiments were repeated 5 times. **(B).** Germination rates on medium supplemented with 120mM NaCl, 0.5 μM MV, 2 μM MV, 100 μM SA or 200 μM Cd. 20 seeds were examined for germination at 3 days/genotype/experiment, and experiments were repeated 5 times. **(C)** Seedling stress survival. 7-day old seedlings were transferred to medium supplemented with 180mM NaCl, 2 μM MV or 400 μM SA and seedling survival examined after 7 days. 10 seedlings were examined per genotype/experiment and experiments were repeated 5 times. Differences in root hair PCD rate, germination and survival for each mutant line was compared to Col-0 using paired t-tests. * ≤ 0.05, ** ≤ 0.01, *** ≤ 0.001, **** ≤ 0.0001. Lines on box and whisker plots display median values while whiskers show the minimum to maximum values.

Additionally, three transgenic lines overexpressing *DTX-1* were obtained from the RIKEN BRC Experimental Plant Division (Ichikawa et al., 2006). Gene expression analysis (qPCR) confirmed overexpression of DTX-1 in *DTX-1 OE2* and *DTX-1 OE3*, but not in *DTX-1 OE1* plants (Figure 6A). Subsequent phenotypic evaluation of the three transgenic lines and wild type plants (Col-0), revealed increased rates of PCD in root hairs of *DTX-1 OE3* 6 hours after HS (Figure 6B). The transgenic lines overexpressing *DTX-1* also showed increased germination under salinity (DTX-1 OE2) and oxidative stress (DTX-1 OE2 and OE3) (Figure 6C), but decreased seedling survival after transfer to medium supplemented with NaCl and MV (DTX-1 OE2 and OE3) (Figure 6D). No significant differences in PCD rates or stress tolerance were observed for *DTX-1 OE1* plants, in line with lack of *DTX1* observed upregulation.

**Figure 6:**
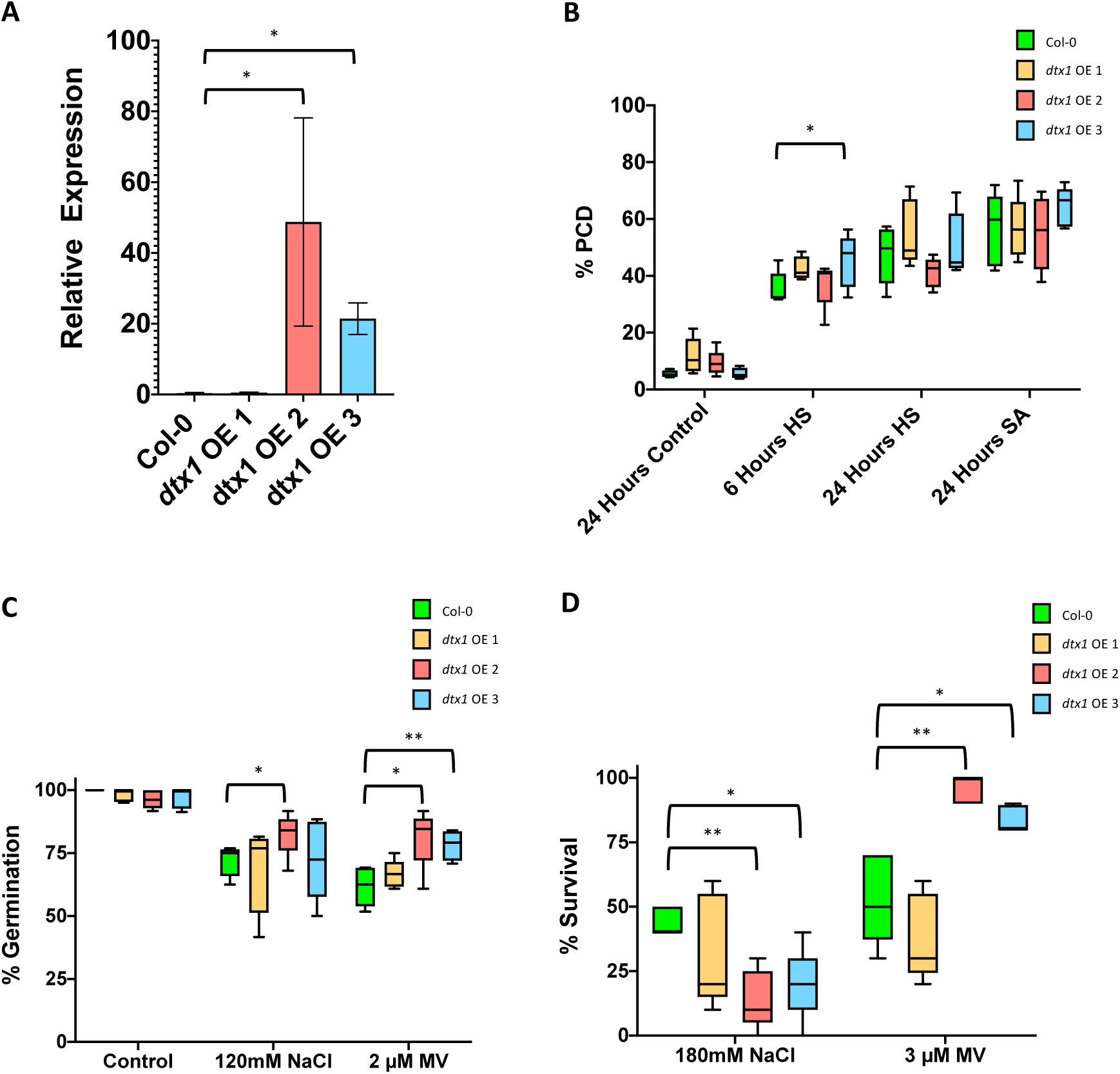
mRNA expression, PCD and stress response phenotypes of DTX1 overexpression lines. (A) mRNA was extracted from 7-day old seedlings and converted to cDNA for qPCR analysis. The presented *dtx1* gene expression was normalized to *actin* and *β-tubulin* (2^-ΔCt^). Indicated p-values were determined using Student’s t-test on log2- transformed relative gene expression data. **(B)** PCD rates in DTX1 overexpression lines. 5 day old seedlings were treated and scored as described previously. Experiments were repeated 5 times. Col-0 was included in each plate and rates of PCD in each mutant line was compared to Col-0. **(C)** Germination rates of DTX1 overexpression lines on medium supplemented with 120mM NaCl and 2 μM MV. Seeds were germinated and scored as described previously. Experiments were repeated 5 times and percentage germination compared to Col-0**. (D)** Seedling survival of the DTX1 overexpression lines. Seedlings were grown and transferred as described previously. Experiments were repeated 5 times and percentage germination compared to Col-0. Differences in root hair PCD rate, germination and survival for each mutant line was compared to Col-0 using paired 1-way ANOVA. * ≤ 0.05, ** ≤ 0.01, *** ≤ 0.001, **** ≤ 0.0001. Lines on box and whisker plots display median values while whiskers show the minimum to maximum values.

### Stress induced transcriptional signatures in *dtx-1* seedlings may explain PCD and stress response phenotypes

To further examine the genetic basis of the PCD and stress sensitivity phenotypes of *dtx1* plants, 7-day old seedlings were subjected to the same mock, HS and SA treatments as those previously used to induce PCD in root hairs. Two hours post treatment whole seedlings were processed for RNA isolation and sequencing. Analysis of reads mapping to the AT2G04040 (DTX1) locus confirmed that the T-DNA insertion was within the CDS and *dtx1* plants were not expressing full transcripts of the DTX1 gene. DGE analysis was performed to identify genes transcriptionally responsive to stress in each line (Col-0, *dtx1*), with adjusted p-value cut off set to 0.05 and fold change cut off set to 5 (DEGs listed in Table S7). This revealed significant overlap, but also notable differences in how *dtx1* and Col-0 seedlings responded to SA and HS treatments, highlighted by genes uniquely up/down regulated in each genotype (Fig 7A). As the similarities and differences between datasets may be more apparent at the pathway level (Zhou et al 2019), the Metascape multiple gene list enrichment analysis tool was used to compare the response to each stress induced in Col-0 and *dtx1* genotypes. The incomplete gene level and functional overlap between the stress response induced in Col-0 and *dtx1* is highlighted by the Circos plots (Fig 7 BC). Across the input gene lists, the enriched terms were clustered into non-redundant groups represented by the most significant term within each cluster (Zhou et al 2019) and enriched clusters unique to each gene list (genotype) are presented in Fig 7 BC, with full list of enriched clusters included in Table S8. Interestingly, many of the identified differences in stress response induced in Col-0 and *dtx1* are linked to processes and pathways previously linked to regulation of PCD in plants. For example, clusters “Response to Ethylene” and “Response to Salicylic Acid”, were enriched in genes downregulated by HS in *dtx1*, but not Col-0 seedlings. Finally, the PlantTFDB TF enrichment tool (Jin et al., 2017) was used to uncover the putative transcriptional regulators underlying the observed response to SA and heat treatments (Table S9). While most of the identified TFs with targets overrepresented among stress-responsive genes in *dtx1* and Col-0 seedlings overlapped, some were genotype specific and had been previously reported to function as regulators of plant PCD in a variety of contexts (Table S9), thus offering a possible explanation for *dtx1* phenotypes. For example, both ANAC017 and ANAC013 were identified as enriched TFs upstream of genes upregulated following HS in *dtx1* plants, but not in Col-0. Similarly, the most significantly enriched TF regulating upregulated ‘core’ PCD genes identified by analyses of cell culture transcriptomes, MYB3R5 (Fig 2E), was enriched for targets among genes upregulated in Col-0 but not in *dtx1* following treatment with HS and SA (Table S9).

**Figure 7.**
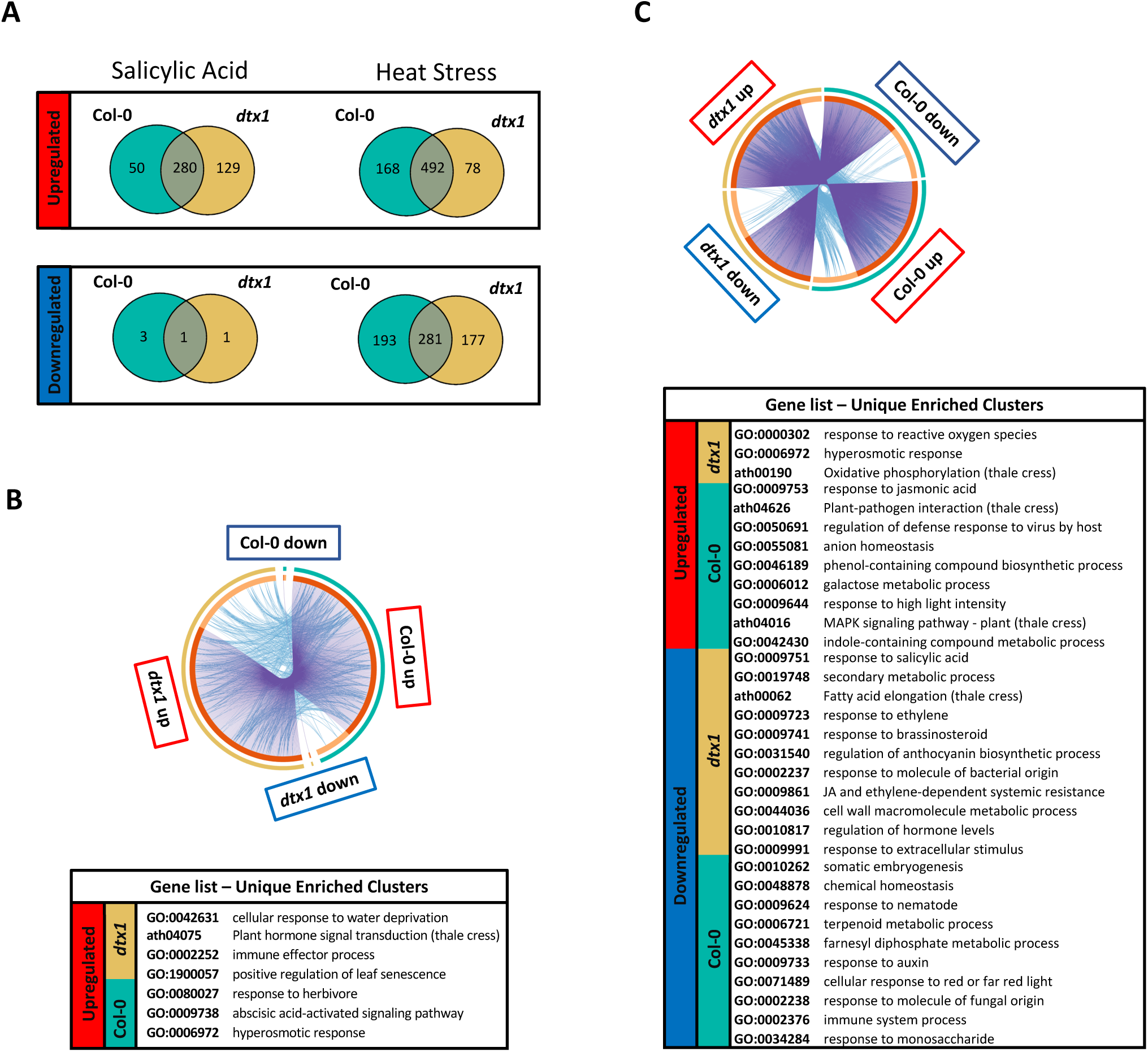
Transcriptional response of *dtx1* and Col-0 seedlings to SA and HS treatments. Venn diagrams present intersection between DEGs up- and down- regulated by applied stress treatments in Col-0 and *dtx1* seedlings **(A)**. Gene level and pathway level overlap in transcriptional response of Col-0 and *dtx1* seedlings to **(B)** salicylic acid and **(C)** heat stress is presented. The external arcs of Circos plots represent the identity of each gene list (Col-0 – teal, *dtx1* – brown). The dark orange segments on the internal arcs represent the genes that appear in multiple lists, and the light orange segments represent genes unique to the gene list. The genes shared by multiple gene lists are linked by purple lines. Blue lines link the genes falling into the same enriched ontology term, indicating functional overlap between the input gene lists. The uniquely enriched clusters for each gene list, represented by the most significant term within each cluster, are presented below each Circos plot.

## Discussion

### Using ACSC and RNA-Seq generates useful resources for the plant PCD research community

The generation of specific transcriptome profiles of well-established systems for investigating PCD is a recommended strategy to reach a thorough understanding of the molecular regulation of this essential process in plants (Olvera-Carrillo et al. 2015). In this study, we draw on the unique advantages of the homogenous ACSC system, which allows PCD induction and modulation using a variety of treatments; and facilitates precise monitoring of the time course of the cell death process. To capture a snapshot of transcriptional changes associated with PCD induction, we sampled the ACSC at time points when PCD rates have started to increase following the stress treatment, but before PCD has been completed in the majority of cells. In this way we hypothesised that the generated snapshot would provide identification of genes that are actively regulated at the initiation of the cell death process.

Differential gene expression analysis revealed only very limited overlap between the transcriptional response induced by salicylic acid (SA, mimic of biotic stress), heat (HS, abiotic stress) and critical dilution (CD, developmental-like cell death) at investigated timepoints, with 10 genes downregulated and 1 gene upregulated by all three stress treatments in Arabidopsis suspension cells (Table S2). This small group of universal stress responsive DEGs included a gene previously suggested to have a pro-PCD function, *AT3G50930*, encoding an Outer Mitochondrial membrane protein of 66 kDa (OM66) (Zhang et al., 2014a), but also *AT1G04400*, encoding a cryptochrome-2, that contributes to plant survival under UV-B stress (Rai et al., 2019). These examples demonstrate that cellular responses to stress can range from activation of pro-survival signalling to initiation of the cell death pathways, and that the ultimate cell fate decision may depend on the balance between these responses (Fulda et al., 2010). Further, this typical simultaneous activation of pro-survival and pro-death responses can hinder isolation of *bona fide* regulators of PCD from transcriptomic datasets.

Here, we attempted to overcome this issue by using each PCD inducer in combination with a PCD inhibitor (SA and 3-MA; HS and LaCl_3_; CD and CM), with the rationale that comparing the transcriptome profiles of cells subjected to stress in absence/presence of PCD inhibitor will allow changes in gene expression related to PCD pathway to be distinguished from the general transcriptional response to stress treatment. Using this approach, we shortlisted three sets of DEGs, and used them to infer regulatory networks underlying three types of cell death programmes: (i) PCD induced by SA and inhibited by 3-MA; (ii) PCD induced by heat and inhibited by LaCl_3_ and (iii) PCD induced by CD and inhibited by CM (Table S3). The generated datasets also enabled selection of ‘core’ PCD genes (i.e. DEGs identified in at least half of all performed comparisons between samples w/wo PCD phenotype: SA vs control, HS vs control, CD vs control, SA vs 3-MA+SA, HS vs HS+LaCl3, CD vs CD+CM; and showing the same direction of regulation), and subsequently, construction of ‘core’ PCD GRN (Fig 4). Our experimental approach and relevance of constructed GRNs in the PCD context was highlighted by their enrichment in terms previously linked to PCD processes in plants. In particular, cluster 1 of the ‘core’ PCD GRN showed enrichment in the terms ‘(Regulation of) Cell Death’ (GO:0010941, GO:008219), but also in “Response to Oxidative Stress” (GO:0006979), “Plant Pathogen Interaction” (ath04626), “Response to Salt Stress” (GO:0009651) and “Cellular Response to Hypoxia” (GO:0071456), referring to conditions well known to induce PCD in plant systems (De Pinto et al., 2012; Van Breusegem and Dat, 2006; Devarenne and Martin, 2007); Shabala, 2009; Fagerstedt, 2010). GO terms enriched by genes from the Cluster 2 of the ‘core’ PCD GRN also refer to processes previously suggested to play a critical role in regulation and execution of plant PCD: cytoskeletal organization (Smertenko and Franklin-Tong, 2011) and cell cycle control (Zebell and Dong, 2015; Kadota et al., 2004). Likewise, many of the TFs with targets overrepresented among identified candidate ‘core’ PCD genes also confirm the relevance of this gene list in the cell death context. For example, the top TF for upregulated ‘core’ PCD candidates, MYB3R5, was previously shown to mediate cell death of stem cells induced by DNA damage (Chen et al 2017). Finally, a known positive regulator of PCD in plants, *metacaspase-5* (Mohamad Zulfazli, 2017), was among the upregulated ‘core’ PCD candidate genes identified.

### Transcriptional changes in a dying cell

The identified ‘core’ PCD candidate genes, their expression pattern and constructed ‘core’ PCD GRN identified three key categories of transcriptional responses associated with early stages of plant PCD: repression of mitochondrial signaling and stress responses, and cell cycle disturbance (Fig 8).

**Figure 8.**
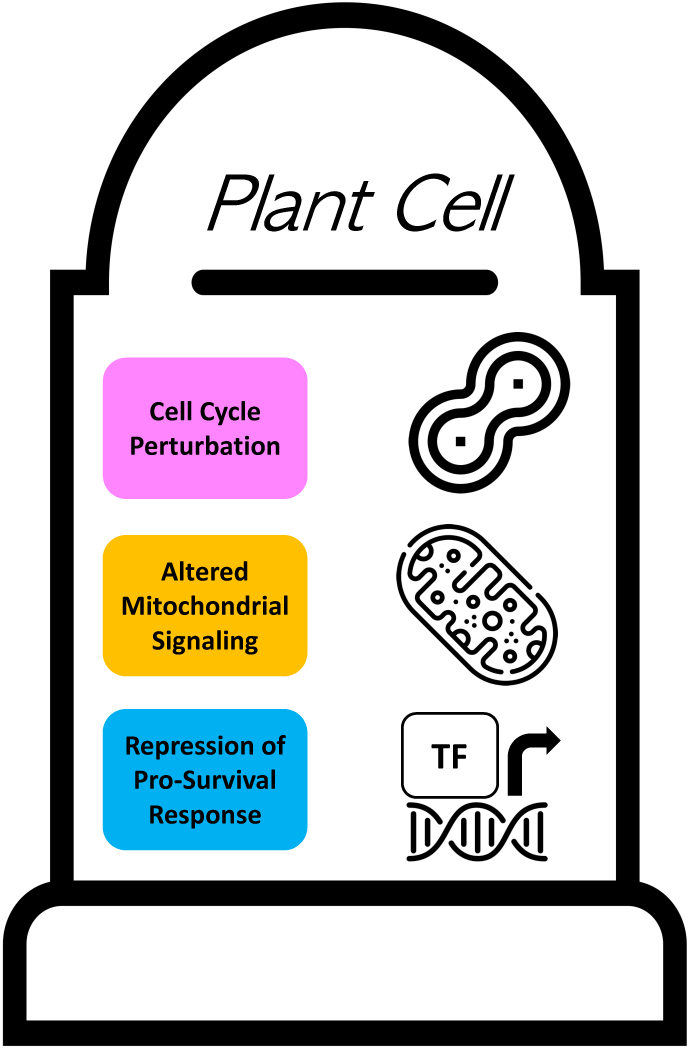
Key transcriptional signatures associated with plant PCD.

#### Repression of Mitochondrial Retrograde Signaling

The ANAC017 and ANAC013 TFs, master regulators of mitochondrial retrograde signaling (Ng et al., 2013; De Clercq et al., 2013; Van Aken et al., 2016) were among downregulated ‘core’ PCD candidate genes. Moreover, mitochondria-nucleus signaling pathway (GO:0031930) was the top enriched GO term among ‘core’ PCD genes (Table S4); and eight other downregulated ‘core’ PCD genes belong to the mitochondrial retrograde signaling regulon (Wang et al., 2018a; Schwarzlander et al., 2012; Van Aken et al., 2009). This includes *Aox1a* that prevents over reduction of the electron transport chain and thus limits the resulting oxidative stress and confers tolerance to multiple abiotic stresses (Vanlerberghe, 2013). These results support the hypothesis that mitochondrial retrograde signalling in plants is at least partly directed towards preventing excessive reactive oxygen species (ROS) formation and thus suppressing cell death (Van Aken and Pogson, 2017).It is therefore plausible that when the cell commits to death, mitochondrial retrograde signalling becomes transcriptionally repressed.

#### Repression of Stress Response

The downregulated ‘core’ PCD genes, represented by Cluster 1 of constructed ‘core’ PCD GRN, include many stress responsive transcriptional regulators with previously suggested pro-survival functions. For example, WRKY25 has been shown to mediate higher tolerance to oxidative stress and delayed senescence (Doll et al., 2020). ZAT10, ZAT12 and SAP12 are zinc finger domain DNA binding and ROS sensitive proteins that have been shown to activate expression of stress response transcripts and thus modulate tolerance to a wide range of abiotic stressors (Mittler et al., 2006; Nguyen et al., 2016; Davletova et al., 2005; Ströher et al., 2009). NAC102 is required for germination under low oxygen stress (Christianson et al., 2009). MYB15 is a positive regulator of resistance to abiotic stress (Ding et al., 2009) and *myb15* cannot contain pathogen triggered hypersensitive cell death (Kim et al., 2020). bZIP60 is involved in the response to ER stress, with homozygous *bzip60* mutants showing markedly weaker induction of many ER stress-responsive genes (Iwata et al., 2008). In addition to these transcriptional regulators, many downregulated ‘core’ PCD genes encode proteins with direct effect on stress tolerance. CML38 is a calcium sensitive protein that localizes to mRNA ribonucleoprotein complexes such as stress granules and enhances resistance to hypoxia (Lokdarshi et al., 2016). AT1G19020 encodes SDA1 that enhances resistance to oxidative stress (Dutta et al., 2020). Further, several downregulated ‘core’ PCD genes identified here, have previously been identified as negative cell death regulators. For example *GILP* (*AT5G13190*) encodes a LITAF domain containing protein that interacts with LSD1 to negatively regulate hypersensitive response associated PCD (He et al., 2011), and copine BON3 (*AT1G08860*) belongs to a family of cell death repressors (Yang et al., 2006). Cluster 1 of the ‘core’ PCD GRN thus represents pro-survival stress response pathways, including both upstream transcriptional regulators and downstream effectors that together enhance tolerance to various forms of abiotic and biotic stress. Following PCD inducing stimuli the almost universal downregulation of the genes in this cluster suggests that one of the core events during PCD in *Arabidopsis* is the early suppression of pro-survival gene transcription. The top predicted transcriptional regulator of the downregulated ‘core’ PCD candidate genes, *AT1G19040* encodes a NAC domain TF that has not yet been characterized in the context of stress response and PCD regulation, and therefore may offer a promising target for future research.

#### Disruption of the Cell Cycle

MYB3R5 was the most significantly enriched TF with targets overrepresented among the upregulated ‘core’ PCD genes, represented by the Cluster 2 of constructed ‘core’ PCD GRN. MYB3R5 was previously shown to mediate cell cycle arrest, growth retardation and PCD of stem cells induced by DNA damage (Chen et al., 2017; Takahashi et al., 2019). The second top TF putatively controlling upregulated ‘core’ PCD genes, MYB119, has also been previously linked to cell cycle regulation, in this case during the F5 stage of female gametogenesis (Rabiger and Drews, 2013). Additionally, Cluster 2 was enriched in genes linked to ‘Mitotic Cell Cycle Process’ (GO:1903047), and Arabidopsis homologues of the anaphase checkpoint control protein MAD2 and cyclin CYCB1;2 (AT5G06150) (Menges et al., 2005) were among upregulated ‘core’ PCD candidate genes. Collectively, this supports perturbation of the cell cycle as a core element of the plant PCD pathway, as previously proposed in effector triggered PCD (Zebell and Dong, 2015).

### Validation of the generated resources in the context of PCD research: discovery of new mediators of plant PCD and stress responses

The reduced complexity of undifferentiated cell suspension cultures is a distinct advantage for studying fundamental biological processes such as PCD (McCabe and Leaver 2000). However, *in vitro* findings may not translate to whole plant systems, especially considering that continuous growth in suspension culture may lead to genomic rearrangements in plant cells (Pucker et al., 2019). To explore the potential of generated datasets to assist isolation of novel genes involved in PCD and stress tolerance, we tested SA and heat stress-induced PCD rates in root hairs of twenty T-DNA mutants using root hair assay (Kacprzyk et al., 2014). Homozygous T-DNA lines in genes encoding putative TFs regulating the PCD process, or hub genes from the constructed GRNs were selected; and 4 lines (*myb119, cmt3, pgip2, dtx1*) displayed consistently reduced PCD levels in root hairs post-treatment compared to wild type control. In addition to the reduced PCD phenotype in root hairs, these T-DNA lines also displayed changes in germination and/or survival under stress conditions compared to wild type plants. *MYB119* encodes a R2R3-MYB class TF with targets overrepresented among identified ‘core’ PCD genes, that was previously linked to female gametogenesis (Rabiger and Drews, 2013). *CMT3*, one of the ‘core’ downregulated genes and a hub node in the SA/3-MA GRN cluster 5, encodes chromomethylase repressing transcription via methylation of non-CG cytosines (Bartee et al., 2001). Although CMT3 has not previously been associated with PCD or with general stress tolerance in plants, there is a growing body of evidence linking changes in chromatin structure and DNA modifications to PCD regulation in plants (Latrasse et al., 2016). *PGIP2*, one of only two upregulated genes in cluster 1 of the HS/LaCl_3_ GRN, encodes a polygalacturonase inhibiting protein expressed as part of the defence response against fungal pathogens (Ferrari et al., 2003). Although expression of *Brassica napus PGIP2* in Arabidopsis (Bashi et al., 2012) and of rice *PGIP2* in *Brassica napus* (Wang et al., 2018b) has been shown to confer increased resistance to the necrotrophic fungus *Sclerotinia sclerotiorum*, this is the first evidence directly linking *PGIP2* to PCD or abiotic stress responses.

Finally, *DTX1* (*<u>D</u>eto<u>x</u>ification <u>1</u>*), selected here as a hub gene in the CD/CM GRN and an element of the ‘core’ PCD GRN, belongs to the multidrug and toxic compound extrusion (MATE) transporters family identified in *Arabidopsis* (Li et al., 2002). The *dtx1* KO mutant showed significant reduction in stress-induced PCD rates and lower germination under oxidative stress, but higher seedling survival under salinity, SA and oxidative (MV) stress conditions. Furthermore, one of two confirmed *DTX1* OE lines showed higher rates of PCD induced by heat after 6h, and both demonstrated increased germination and survival under MV treatment compared to wild type. The transcriptional response of *dtx1* and Col-0 seedlings treated with heat and SA stress was also distinctly different. In contrast to transcriptomes of homogenous suspension cells undergoing PCD, the whole seedling transcriptomes represent changes in gene expression that occur in a whole range of cell types and tissues, with many of them activating processes preceding or counteracting cell death, which makes it difficult to interpret the observed differences between *dtx1* and Col-0 in the context of PCD. However, downstream targets of MYB3R5, TF known to promote cell death induced by DNA damage (Chen et al 2017), here identified as a putative upstream transcriptional regulator of the upregulated ‘core’ PCD genes, were overrepresented among both heat stress and SA-induced genes in Col-0, but not in *dtx1* plants, offering a potential explanation for observed phenotypic differences between them. While these results fail to clearly identify *DTX1* as a pro-survival or pro-PCD mediator, potentially due to functional redundancy with other DTX family members, they highlight its relevance in the context of cell death and stress tolerance. This is in line with numerous DTX/MATE proteins being previously linked to various aspects of biotic and abiotic stress resistance. For example, DTX50 functions as an ABA efflux transporter and regulates drought tolerance (Zhang et al., 2014b) and a gain of function mutation in *Arabidopsis* DTX6 has been shown to confer resistance to MV (Xia et al., 2021), similar to observations for DTX1 presented here. Further, the overexpression of a DTX family gene from cotton has also been shown to confer widespread abiotic stress tolerance in *Arabidopsis* (Lu et al., 2019). *EDS5* (*DTX47*) encodes a MATE transporter that localizes to the chloroplasts envelope where it exports SA to the cytoplasm (Serrano et al., 2013; Yamasaki et al., 2013), and is thus essential for the activation of downstream SA signalling pathway and regulating disease resistance (Nawrath et al., 2002). Further studies on the role of DTX1 in PCD and stress responses in plants are therefore well warranted.

In conclusion, our results indicate that our approach based on cell suspension culture, a well-established model for studying plant PCD, and RNA-Seq was successful in isolation of promising candidate PCD genes and their transcriptional regulators. Combining three well characterized PCD inducers, chosen to represent different contexts in which PCD occurs in plants, with three different PCD inhibitors yielded new insights into regulatory networks underlying both ‘core’ and stimuli-specific cell death programmes. The constructed GRNs not only shed light on how PCD is regulated, but also facilitate identification of hub signaling genes, that are highly connected and thus likely to play a role in gene regulation of biological processes (Yu et al., 2017). The generated resources are therefore of interest to the plant PCD community and should accelerate research in this area.

## Materials and Methods

### ACSC growth conditions and treatments

The ACSC (Ler ecotype) were maintained as previously described (May and Leaver, 1993) and subcultured every 7 days by taking 10 mL of culture and adding it to 100 mL of fresh medium. Cells grown in the dark at room temperature (21°C) were used for experiments. Salicylic acid (SA)/3-methyladenine (3-MA) experiments: 6-day old cells (20 mL in 100 mL flask) were treated with 3-MA (ChemCruz) (15 mM) or solvent control (sterile distilled water, SDW) and returned to standard growth conditions. The 0.1 M 3-MA stock in SDW had to be re-dissolved by heating prior to use (Kacprzyk et al., 2014). After 24 hours, the cells were treated with SA (Merck) (3.25 mM) or solvent control (0.65% EtOH). Flasks were again returned to standard growth conditions until monitoring of cell death/sampling for total RNA extraction at indicated time points.

Heat-stress (HS)/LaCl_3_ experiments: 6-day old cells (20 mL in 100 mL flask) were treated with LaCl_3_ (Thermo Scientific) (750 µM) or solvent control (SDW) 10 minutes prior to heat treatment. Heat treatment (54°C, 10 minutes) was performed in Grant OLS200 waterbath with shaking (100 rpm). Following heat treatment, flasks were returned to standard growth conditions until monitoring of cell death/sampling for total RNA extraction at indicated time points.

Critical dilution (CD)/conditioned medium (CM) experiments: CM was prepared from 6-day old light grown ACSC. Briefly, 100 mL of 6-day old suspension cells was filtered through a 75 μm nylon mesh, filtered under vacuum through Whatman grade 1 paper using a Buchner funnel. The resulting filtrate (approximately 70-80 mL) was supplemented with NAA (Duchefa Biochemie) (0.05 mg/L) and kinetin (Duchefa Biochemie) (0.05 mg/L) to replenish phytohormones as described in (Daly, 2013).

The pH was adjusted to 5.8 and resulting CM autoclaved at 120°C for 20 minutes. CM was stored at 4°C and used within 24 hours. To induce PCD via dilution below critical cell density, 7-day old dark grown cultures were sub-cultured at 10x dilution (control) or 50x dilution (below critical density) into fresh culture medium, or fresh medium supplemented with 20% CM in a total volume of 20 mL. Samples were then returned to standard growth conditions until monitoring of cell death/sampling for total RNA extraction at indicated time points.

### Arabidopsis plant material and growth conditions

Arabidopsis seedlings were grown aseptically as described previously (Hogg et al., 2011). All mutant and transgenic lines, obtained from the Nottingham Arabidopsis Stock Centre (Alonso José et al., 2003, Scholl et al., 2000) and RIKEN BRC Experimental Plant Division (Ichikawa et al., 2006) used in this study are detailed in Table S6. The used Arabidopsis FOX lines overexpressing *DTX1* were developed by the plant genome project of RIKEN Plant Science Center. The Salk mutants showing PCD phenotypes were genotyped using primers listed in Table S10 to confirm homozygous T-DNA (transfer DNA) insertion.

### Induction of PCD in Arabidopsis root hairs

Arabidopsis seedlings were grown aseptically on semi-solid half-strength Murashige and Skoog medium (MS basal salts, 2.15 g L^-1^, 1% sucrose, pH adjusted to 5.8, solidified with 1% agar) in 12 cm square Petri dishes placed in vertical position at standard growth conditions (22°C, 16 hr light/8 hr dark). Five-day old seedlings were used for experiments. For PCD induction using heat, seedlings were transferred to a 24-multiwell plate, with each well containing 1 mL of SDW. Plates were then floated in a Grant OLS200 water bath (no mixing) at 45°C for 10 minutes. For PCD induction using SA, seedlings were placed in 30 μM SA solution in SDW in 24-multiwell plate. Following the treatments, plates with seedlings were returned to growth conditions until scoring for PCD, necrosis and viability rates.

### Cell Death Assay

The levels of PCD, necrosis and viability in suspension cells/root hair cells were determined as previously described (Kacprzyk et al 2017; Hogg et al 2011). Briefly, cells/seedlings were stained in a 1 μg/ml solution of FDA on the standard microscope slides and immediately examined under white light and blue light.

Cells/root hairs positive for FDA staining were scored as alive, whereas cells/root hairs negative for FDA staining were scored as PCD if they exhibited protoplast retraction away from the cell wall, or as necrotic when this hallmark morphology was absent. For ACSC, at least 200 cells were scored per sample and for the Arabidopsis seedlings, 2 two seedlings were scored per genotype per condition (approximately 300 root hairs examined) during each experimental repeat.

### RNA Extraction, Quantification, RNA-Seq and Quantitative PCR

Harvesting of material: (1) *ACSC:* At indicated time point, 10 mL of cell suspension were removed from each flask and media was removed via vacuum filtration using a Buchner funnel and Whatmann grade 1 paper. The cells were then transferred to 2 mL round bottom Eppendorf tubes prefilled with approximately 20 chrome-steel beads, flash frozen using liquid N_2_ and stored at – 80°C until RNA extraction. (2) Arabidopsis seedlings: 2 hours after treatment, ten seedlings per treatment per genotype were removed from the multiwell plates, dried on tissue paper, transferred to 2 mL round bottom Eppendorf tubes and frozen in liquid N_2_ as above. For total RNA isolation, plant material was homogenized in a mixer mill (Retsch MM400), with 4 minutes of horizontal oscillation (20 oscillations/s). The QIAGEN RNeasy Plant Mini kit was used according to the manufacturer’s instructions, including the on-column DNAse treatment step. The resulting total RNA was quantified using Nanodrop (Thermo Scientific) and RNA quality was confirmed using Bioanalyzer (Agilent).The library preparation and mRNA sequencing (poly-A selection, paired-end 150bp reads, 6G clean data per sample) was performed by Novogene (Cambridge, UK).

### qPCR analysis

cDNA was synthesized using ThermoScientific first-strand cDNA synthesis kit according to the manufacturer’s instructions. Real-time PCR was carried out on Applied Biosystems ViiA 7 thermocycler using KAPA SYBR® FAST Universal master mix according to manufacturer’s instructions. Reaction conditions were: 3 min 95°C, 40 cycles: 5s at 95°C, 30s at 58°C with melting curve analysis included. Primers used (5’ è 3’): *dtx-1* - Forward GATACGCATTCAGCAACAGC, Reverse ATGTGTTGCCAACCACTTCCT; *actin* - Forward TCCGTTTTGAATCTTCCTCAATCTCA, Reverse TTGAATATCATCAGCCTCAGCCATT; *β-tubulin* - Forward: CTCAAGAGGTTCTCAGCAGTA Reverse: TCACCTTCTTCATCCGCAGTT (Kim et al. 2007). *DTX1* transcript levels were normalized to *actin* and *β-tubulin* for each genotype using geometric mean of their expression (Vandesompele et al., 2002).

### Differential Gene Expression (DGE) analysis

The DGE analysis was performed using Galaxy’s graphical user interface (Afgan et al., 2018). The FASTQ files were uploaded to Galaxy platform and the quality of raw reads was first assessed using FastQC (Andrews, 2010) using default settings.

Paired reads were then trimmed using Trimmomatic (Bolger et al., 2014), using the sliding window function to trim windows of 4 bases that dropped below an average Phred score of 20 (99% accuracy). Trimmed reads were mapped to the TAIR10 *Arabidopsis* genome (Lamesch et al., 2012) using HISAT2 (Kim et al., 2019), with strand information set as ‘unstranded’. Mapping yielded an overall alignment rate of ∼94 – 96%. The expression value counts for the aligned reads were assigned using the htseq-count tool in HTSeq (Anders et al., 2015) in union mode, with a minimum alignment quality set to ten. Subsequently, differential gene expression analysis was carried out using the DESeq2 package (Love et al., 2014), with parametric fitting and with independent filtering enabled. The Benjamini and Hochberg (Benjamini and Hochberg, 1995) method was used for p-value adjustment, with adjusted p-value cut-off applied 0.05. Results of DESeq2 analysis were then imported into R studio version 2022.2.3.492 (RStudio Team, 2020) for data handling and visualization.

### Transcription Factor Enrichment Analysis

The TF enrichment tool of the Plant Transcriptional Regulatory Map (PlantRegMap) (Jin et al., 2017) was used to identify TFs with overrepresented targets for the upregulated and downregulated candidate PCD genes using the motif method and 0.05 adjusted p-value cut-off.

### Metascape Gene Annotation and Analysis

The Metascape online platform (Zhou et al., 2019) was used for functional enrichment analysis using default parameters. For DEGs identified following treatment with PCD blockers 3-MA and LaCl_3_ a single gene list analysis was performed, and for comparisons of DEGs in Col-0 and *dtx1* plants following treatment with HS and SA the multiple gene list Metascape function was employed.

### Regulatory Network Inference

The lists of candidate genes for regulators of stimuli specific- and ‘core’- PCD response were used for inference of the underlying regulatory networks. For HS- and CD-induced PCD, as well as for the ‘core’ PCD candidate gene lists (including 59, 16 and 401 DEGs respectively), the GeneMANIA app (Warde-Farley et al., 2010) was used in Cytoscape (Shannon et al., 2003) for construction of gene regulatory network (GRN). Subsequently, Clustermaker (Morris et al., 2011) was used to carry out MCL (Markov Cluster Algorithm) clustering with granularity set to 2.5. The genes in each cluster where then analysed for Gene Ontology enrichment using Metascape, and the significantly enriched categories (GO Biological Processes and KEGG Pathways) overlayed over the relevant genes in each cluster. Due to the already large number (1173) of candidate genes identified for SA-induced PCD, rather than GeneMANIA, the STRING App (Doncheva et al., 2019) was instead used to generate a GRN consisting only of high confidence protein level interactions (using the protein query database, with a confidence score cut-off of 0.7 and no additional interactors allowed). The resulting network was then clustered using STRINGs inbuilt MCL clustering program, with a granularity setting of 2.5. The six largest clusters (those with >11 nodes) were selected and the genes in each analysed for Gene Ontology enrichment using Metascape.

### Stress Germination Assay

Wild type (Col-0) and mutant/transgenic seeds (20 seeds per line) were germinated on semi-solid half-strength Murashige and Skoog medium (MS basal salts, 2.15 g L^-1^; 1% sucrose; 0.6% agar, pH 5.8), supplemented with 120mM NaCl (Duchefa Biochemie), 0.5μM/2μM methyl viologen (MV) (ThermoScientific), 100 μM SA (Merck), or 200 μM cadmium chloride (CdCl_2_) (Sigma-Aldrich). Seeds were stratified for two days at 4°C in the dark before transfer to growth conditions (22°C, 16h light/8h dark). Germination after 3 days (22°C, 16h light/8h dark) was assessed using a dissecting microscope. Experiments were repeated 5 times.

### Stress Survival Assay

Seven-day old seedlings were grown on semi-solid half-strength Murashige and Skoog medium (MS basal salts, 2.15 g L^-1^; 1% sucrose; 0.6% agar, pH 5.8) at 22°C, 16h light/8h dark, before being aseptically transferred (10 seedlings per line) to the same medium supplemented with 180mM NaCl, 2 μM MV or 400 μM SA and grown under the same conditions for 7 days. Seedling survival was scored based on the bleaching of the newest emerging leaves.

### Statistical Analysis

Statistical analysis of *Arabidopsis* seedlings was carried out using GraphPad Prism version 8.2.1 for Mac OS (GraphPad Software, San Diego, California USA).To account for biological variation during different experimental replicates, Col-0 plants were included alongside the relevant t-DNA or overexpressor mutants for every experiment. The response of each t-DNA or overexpressor mutant line was then compared to the Col-0 plants from the same experiments using paired t-tests or paired one-way anova tests (without false discovery rate correction) for t-DNA or overexpressor lines respectively.

### Accession Numbers

The RNA-Seq data used in this publication has been uploaded to the NCBI Sequence Read Archive and is available under the Bioproject Accession number PRJNA926379. The AGI identifiers for all genes mentioned in the manuscript are listed in table S11.

## Supplemental Figures

**Figure S1.**
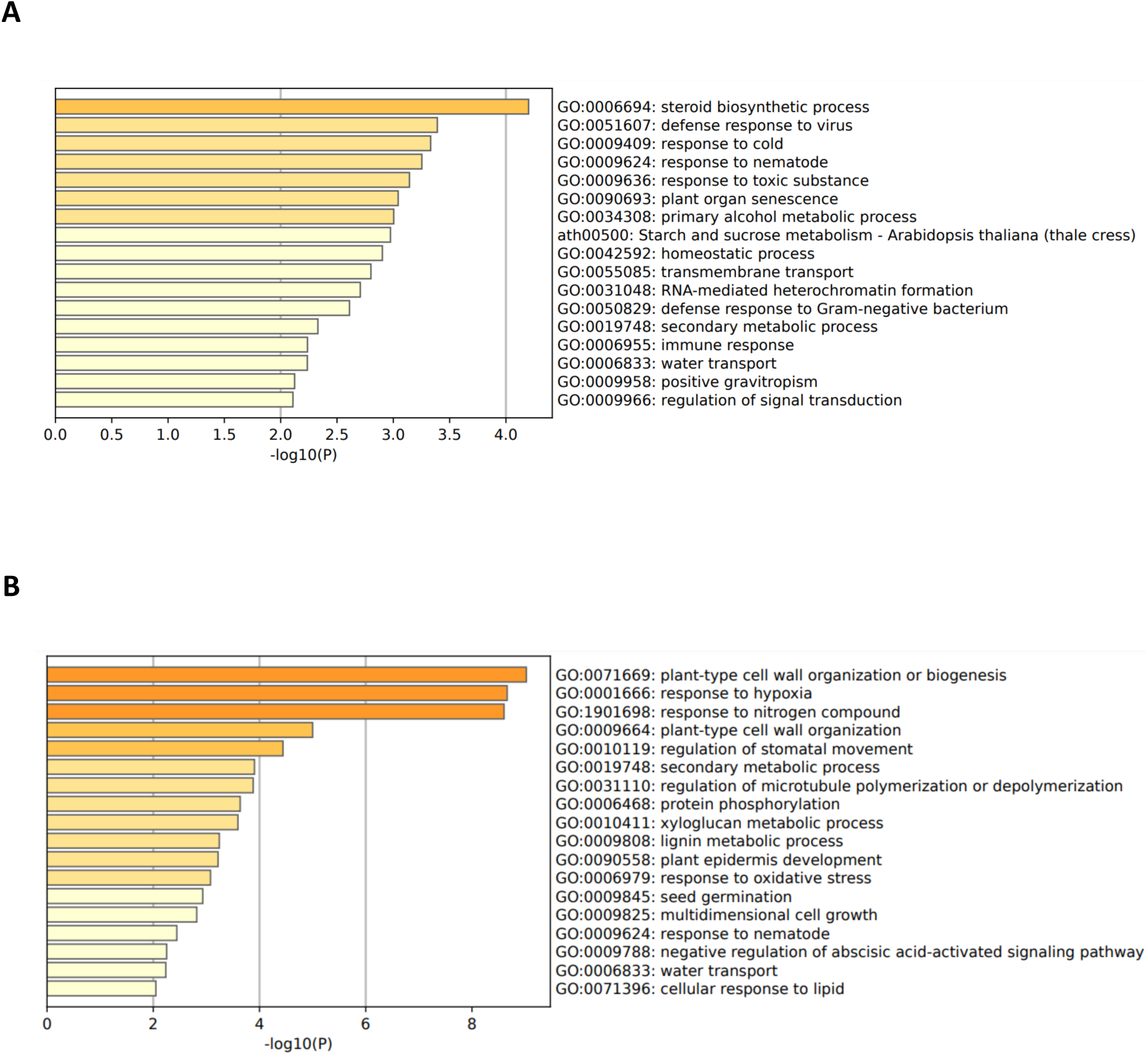
Transcriptional response of ACSC to 3-MA and LaCl_3._ Significantly enriched clusters of GO terms for DEGs in response to treatment with 3-MA (A) or LaCl_3_(B).

**Figure S2.**
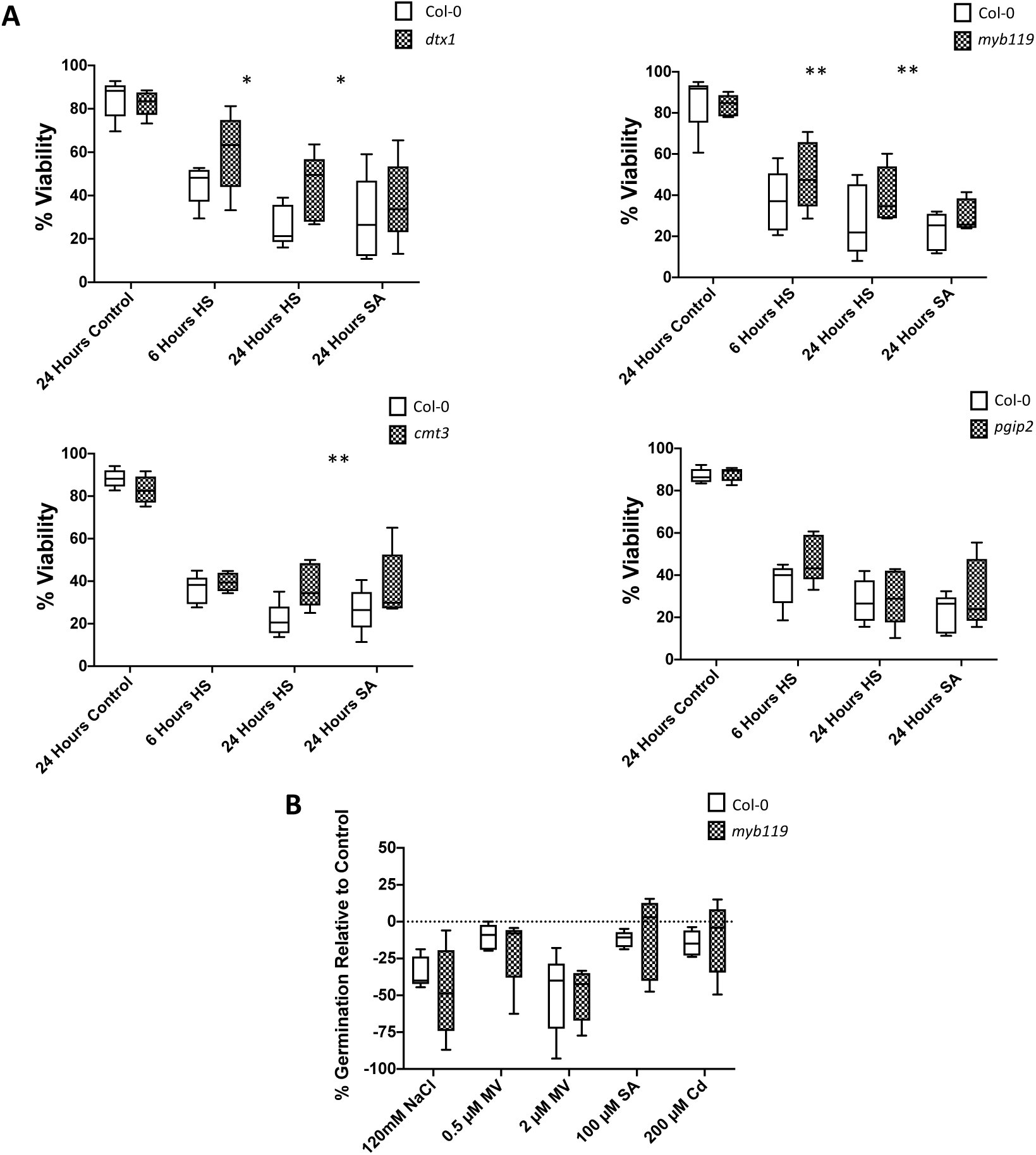
Viability and germination rates of T-DNA insertion mutants. (A) 5 day old seedlings were transferred to a 24 well plate, with each well containing 1 mL of SDW. Seedlings were then subjected to either a 10-minute HS at 45°C or treated with 30 μM SA and scored after 6 and 24 hours or 24 hours respectively. For each time point, the root hairs of 2 seedlings of each line were scored as viable, PCD or necrotic. Experiments were repeated 5 times. **(B)** Germination rates of *myb119* plants on medium supplemented with 120mM NaCl, 0.5 μM MV, 2 μM MV, 100 μM SA or 200 μM Cd. 20 seeds were examined for germination at 3 days/genotype/experiment, and experiments were repeated 5 times. For each experiment, the change in germination relative to control conditions was calculated to account for reduced germination rates in *myb119* plants. Differences in root hair viability rate and germination for each mutant line was compared to Col-0 using paired t-tests. * ≤ 0.05, ** ≤ 0.01, *** ≤ 0.001, **** ≤ 0.0001. Lines on box and whisker plots display median values while whiskers show the minimum to maximum values.

## Supplemental Data

Figure S1. Enrichment analysis of genes responsive to 3-MA and LaCl_3._

Figure S2. Viability of T-DNA insertion lines displaying altered PCD rates during the root hair assay.

Table S1. Full RNA-Seq DESeq2 Cell Culture Results

Table S2. List of 11 DEGs in common between all three PCD inducers and control samples.

Table S3. List of candidate genes involved in regulation of PCD induced by each stress stimuli (SA, HS, CD).

Table S4. List of ‘core’ PCD genes.

Table S5. TFDB results for shortlisted stress stimuli- and ‘core’- PCD candidate genes.

Table S6. Genes and corresponding T-DNA mutants selected for phenotyping. Table S7. DEG lists for Col-0 and *dtx1* plants treated with HS and SA.

Table S8. Metascape gene ontology enrichment results for Col-0 and *dtx1* plants treated with HS and SA.

Table S9. TFDB results for Col-0 and *dtx1* plants treated with HS and SA. Table S10. List of primers used for genotyping of T-DNA mutants.

Table S11. Accession numbers of all mentioned genes.

